# Discovery of a Selective, State-Independent Inhibitor of Na_V_1.7 by Modification of Guanidinium Toxins

**DOI:** 10.1101/869206

**Authors:** H Pajouhesh, JT Beckley, A Delwig, HS Hajare, G Luu, D Monteleone, X Zhou, J Ligutti, S Amagasu, BD Moyer, D Yeomans, J Du Bois, JV Mulcahy

**Affiliations:** SiteOne Therapeutics, South San Francisco, CA 94080; SiteOne Therapeutics, Bozeman, MT 59718; Department of Chemistry, Stanford University, Stanford, CA 94305; Neuroscience Department, Amgen Research, Thousand Oaks, CA 91320; Department of Anesthesiology, Perioperative, and Pain Medicine, Stanford University, Stanford, CA 94305

## Abstract

The voltage-gated sodium channel isoform Na_V_1.7 is highly expressed in small diameter dorsal root ganglion neurons and is obligatory for nociceptive signal transmission. Genetic gain-of-function and loss-of-function Na_V_1.7 mutations have been identified in select individuals, and are associated with episodic extreme pain disorders and insensitivity to pain, respectively. These findings implicate Na_V_1.7 as a key pharmacotherapeutic target for the treatment of pain. While several small molecules targeting Na_V_1.7 have been advanced to clinical development, no Na_V_1.7-selective compound has shown convincing efficacy in clinical pain applications. Here we describe the discovery and characterization of ST-2262, a Na_V_1.7 inhibitor that blocks the extracellular vestibule of the channel with an IC_50_ of 72 nM and greater than 200-fold selectivity over off-target sodium channel isoforms, Na_V_1.1–1.6 and Na_V_1.8. In contrast to other Na_V_1.7 inhibitors that preferentially inhibit the inactivated state of the channel, ST-2262 is equipotent against resting and inactivated protein conformers. In a non-human primate model, animals treated with ST-2262 exhibit markedly reduced sensitivity to noxious heat. These findings establish the extracellular vestibule of the sodium channel as a viable receptor site for selective ligand design and provide insight into the pharmacology of state-independent inhibition of Na_V_1.7.

**Significance Statement:** Pain is among the most common reasons for seeking medical care, yet many frequently prescribed drugs, particularly the opioids, cause problematic side effects and carry a risk of addiction. Voltage-gated sodium ion channels (Na_V_s) have emerged as promising targets for the development of non-opioid pain medicines. Na_V_s are involved in the propagation of electrical signals along neurons throughout the body. Humans born without a functional copy of one sodium channel subtype, Na_V_1.7, are unable to experience most types of pain. In the present work, we disclose the discovery and characterization of a selective inhibitor of Na_V_1.7 that reduces sensitivity to a painful thermal stimulus in non-human primates. Findings from this work may help guide the development of novel, non-addictive drug candidates as alternatives to opioids.

## Introduction

The voltage-gated Na^+^ ion channel (Na_V_) isoform 1.7 has emerged as a high-interest target for the discovery of non-opioid pain therapeutics based on its preferential expression in small diameter peripheral sensory neurons and compelling validation from human genetics.(1) Na_V_1.7 loss-of-function mutations result in whole-body insensitivity to pain; conversely, gain-of-function variants are associated with episodic extreme pain disorders and small fiber neuropathies.(2–5) Discovery of selective inhibitors of Na_V_1.7 has been challenging due to the structural conservation of off-target Na_V_ isoforms (Na_V_1.1–1.6, Na_V_1.8 and Na_V_1.9), inhibition of which is likely to result in safety liabilities.(6–8)

Na_V_s are integral membrane proteins expressed in excitable cells that comprise a ~260 kD pore-forming α-subunit and two accessory β-subunits (Figure 1).(9) The central pore of the α-subunit is encircled by four voltage-sensing domains (VSD I–IV). Channel gating occurs through protein conformational changes in response to membrane depolarization. At least nine discrete binding sites on the Na_V_ α-subunit have been identified for peptides and small molecules that influence ion conductance.(10) The large majority of molecules that engage Na_V_s bind preferentially to a specific conformational state of the channel and show use-dependent activity. Clinical Na_V_ inhibitors (e.g., bupivacaine, lidocaine, carbamazepine) are both state- and frequencydependent agents that lodge in the intracellular pore of the α-subunit, a site that is highly conserved between isoforms. These drugs rely on local administration or frequency dependent inhibition to achieve a margin between the desired pharmacodynamic effect and dose-limiting side effects. Investigational Na_V_ inhibitors, such as peptide toxins isolated from venomous species, interact with VSDs, trapping the S4 helix in either an activated or deactivated conformation.(11, 12) Similarly, a class of small molecule aryl and acyl sulfonamide compounds bind to the activated conformation of VSD IV (Figure 1b).(13–16) By contrast, cationic guanidinium toxins and peptide cone snail toxins inhibit ion conduction by sterically occluding the extracellular vestibule of the channel pore (Site 1). The former are a unique collection of natural products exemplified by saxitoxin and tetrodotoxin – high affinity, state independent blockers against six of nine Na_V_ subtypes (Figure 1c).(17)

**Figure 1.**
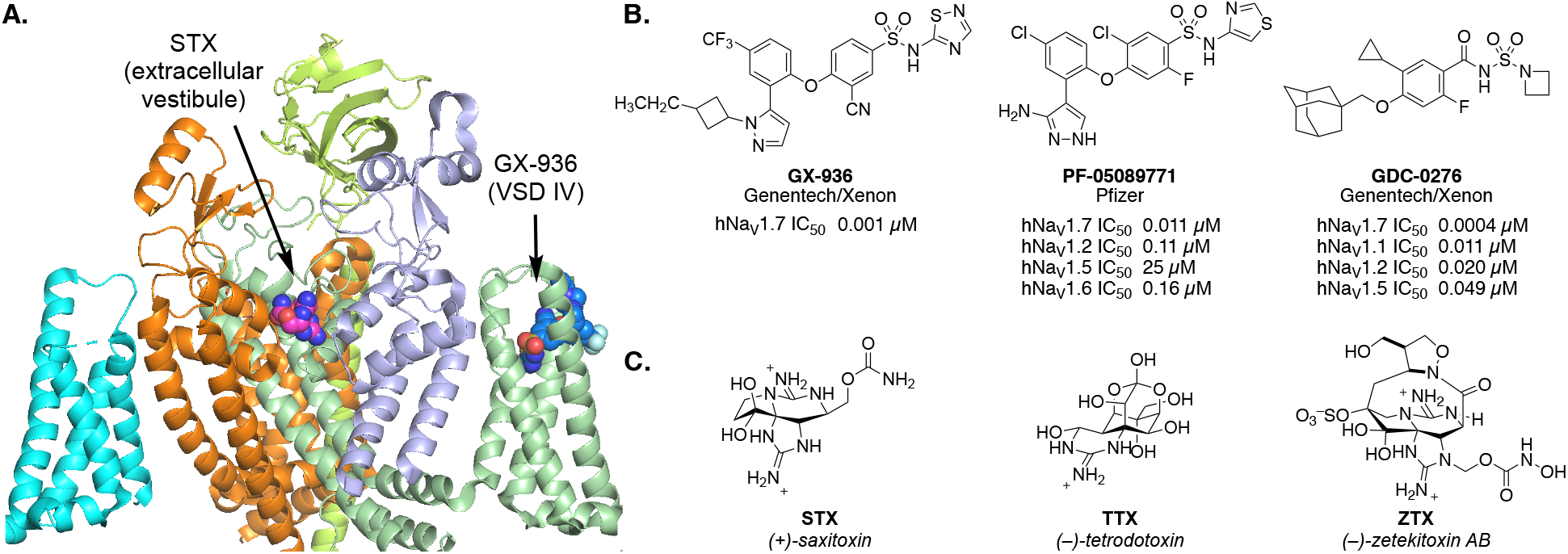
**(a)** Cryo-EM structure of STX bound to human Na_V_1.7-β1-β2 complex (PDB: 6j8g) with GX-936 positioned approximately based on PDB: 5ek0. **(b)** Representative Na_V_1.7 inhibitors that bind VSD IV. **(c)** Natural Na_V_ inhibitors that bind to the extracellular vestibule.(18–21)

In the pursuit of isoform-selective inhibitors of Na_V_1.7, two binding sites, the cystine knot toxin site at VSD II and the sulfonamide site at VSD IV, have been heavily interrogated. Certain cystine knot toxins that engage VSD II such as HwTx-IV, Pn3a and ProTx-II exhibit 6–1000x selectivity for Na_V_1.7 over other channel isoforms. Potency and selectivity for this target have been improved with synthetic toxin derivatives.(22–26) Among small, Lipinski-type molecules, only the aryl and acyl sulfonamides pioneered by Icagen/Pfizer and subsequently investigated by Amgen, Chromocell, Genentech/Xenon, Merck and others have shown evidence of significant Na_V_1.7 isoform selectivity.(16, 7) Within the sulfonamide series, selectivity levels are >1000x over certain off-target isoforms including the cardiac isoform, Na_V_1.5, but generally 10–50x against Na_V_1.2 and Na_V_1.6. Many but not all sulfonamide Na_V_1.7 inhibitors suffer from undesirable pharmaceutical properties, including high plasma protein binding (e.g. >99.8%), cytochrome p450 inhibition, *in vitro* hepatotoxicity and high unbound clearance,(27, 28) which have hindered clinical development. Although a number of candidates have been advanced to human testing, one compound has been discontinued after a Phase 2 study likely due to limited efficacy (PF-05089771); others have been discontinued after Phase 1 trials for reasons that may be related to safety liabilities such as elevated expression of liver transaminases and hypotension (GDC-0276).(29, 30)

Electrophysiology studies with naturally occurring Site 1 ligands against different wildtype and mutant Na_V_ isoforms have identified the extracellular vestibule of Na_V_1.7 as a promising locus for selective inhibitor design.(31–33) The outer mouth of the channel is formed from residues that link the S5–S6 helices (referred to as pore loops) from each of the four domains. The domain III pore loop of human Na_V_1.7 contains a sequence variation, 1398T/1399I, that is not present in other human Na_V_ subtypes (which contain MD at equivalent position, Suppl Table 1).(31) Comparison of the amino acid sequence of the domain III pore loop across species indicates that this variation is unique to primates. The half maximal inhibitory concentration (IC_50_) value for saxitoxin (STX) is markedly altered (250-fold change) depending on the presence or absence of the 1398T/1399I variant. Against rNa_V_1.4, the IC_50_ of STX is 2.8 ± 0.1 nM compared to hNa_V_1.7 for which the IC_50_ = 702 ± 53 nM.(31) Introduction of the alternative domain III pore loop sequence by mutagenesis restores potency (hNa_V_1.7 T1398M/I1399D IC_50_ = 2.3 ± 0.2 nM). These findings suggest that it may be possible to capitalize on structural differences in the extracellular vestibule between hNa_V_ isoforms to design Na_V_1.7-selective inhibitors.

Recent advances in the *de novo* synthesis of guanidinium toxin analogues have enabled systemic examination of the structure-activity relationship (SAR) properties that govern hNa_V_1.7 potency and isoform selectivity.(34–37) Prior to 2016, the binding orientation of STX proposed in the literature indicated that the C11 methylene carbon was positioned proximally to domain III pore loop residues.(38–40) SAR and mutant cycle analysis studies posited a revised pose in which the C13 carbamate moiety abuts DIII.(32) This revised binding pose was recently confirmed by cryoelectron microscopy (cryo-EM) structures of saxitoxin bound to Na_V_Pas and hNa_V_1.7.(18, 41) In the present study, analogues of STX substituted at both the C11 and C13 positions were investigated to understand the requirements for selective inhibition of hNa_V_1.7. These efforts led to the discovery of ST-2262, a potent and selective inhibitor of Na_V_1.7 that is efficacious in a non-human primate (NHP) model of noxious thermal sensitivity.

## Results

### Discovery of ST-2262

ST-2262 was discovered through a rational design strategy aimed at identifying analogues of natural bis-guanidinium toxins that preferentially inhibit hNa_V_1.7 over other off-target hNa_V_ isoforms.(31) Mutagenesis, homology modeling, and docking studies conducted prior to 2016 suggested that bis-guanidinium toxins orient in the outer mouth of the channel with the C11 methylene center positioned toward the domain III pore loop of Na_V_ (Figure 2a, Original pose).(38–40) Exploration of substitution at C11 of decarbamoyl saxitoxin (dcSTX) led to the identification of a series of analogues bearing aryl amide groups at this site. Certain compounds, as exemplified by ST-282, show excellent potency against hNa_V_1.7 but minimal selectivity (~1:1) over off-target isoforms such as hNa_V_1.4 (Figure 2b). The finding that hNa_V_1.7 isoform selectivity could not be achieved by modification of the C11 substituent led us to investigate SAR at alternative positions. These studies followed evidence that the proper binding orientation of STX is rotated ~180° from earlier models, thus placing the C13 substituent in close proximity to domain III (Figure 2a, Revised pose).(32)

**Figure 2.**
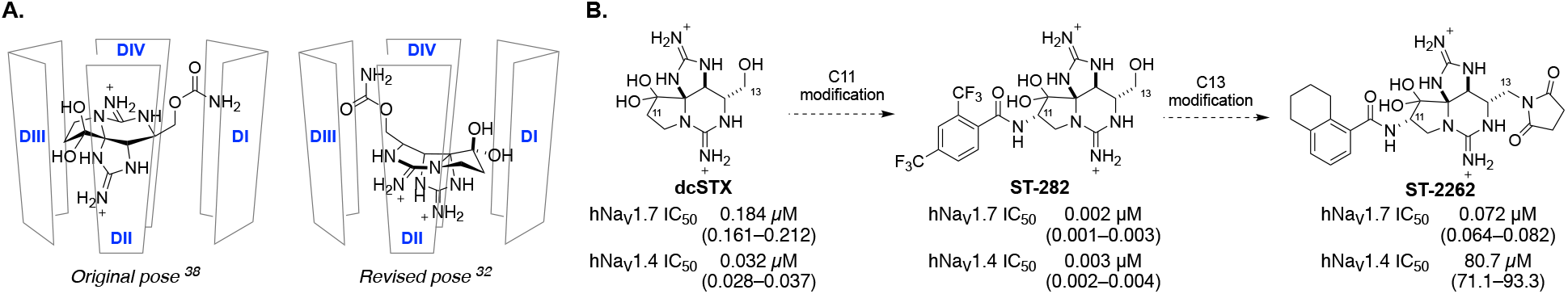
**(a)** The consensus pose for binding of STX in the extracellular vestibule of Na_V_ oriented C11 in proximity to the DIII pore loop prior to 2016.(38) A revised pose based on mutant cycle analysis and recent cryo-EM structures orients the C13 carbamate near DIII.(32, 41) **(b)** ST-2262 was discovered by a rational design strategy that aimed to identify functional groups that interact with the DIII 1398T/1399I sequence variation unique to primate Na_V_1.7. Values are mean (95% CI).

Analogues of STX bearing amides, carbamates, esters, ethers and urethanes at the C13 position were prepared in an effort to identify compounds with improved selectivity for hNa_V_1.7. Insight from studies of a naturally occurring STX C13 acetate congener, STX-OAc, helped guide selection of compounds for synthesis (Suppl Figure 1).(32) The difference in potencies between STX and STX-OAc is striking considering that their structures vary at a single position (NH_2_→CH_3_). Following this lead, we explored substituents at C13 that could replace the hydrolytically unstable acetate group. Ultimately, the C13 succinimide was discovered as a suitable acetate isostere, which was paired with a C11 tetrahydronaphthyl amide to generate ST-2262, the focus of the present study.

### ST-2262 is a potent, selective, state-independent inhibitor of hNa_V_1.7

The potency of ST-2262 against hNa_V_1.7 stably expressed in HEK293 cells was assessed by manual patch clamp electrophysiology with a voltage-protocol that favors the resting state of the channel. Using a voltage protocol with a holding potential of –110 mV and a stimulus frequency of 0.33 Hz, the IC_50_ of ST-2262 against hNa_V_1.7 was measured at 0.072 μM (95% confidence interval (CI) 0.064–0.082) (Figure 3a, Suppl Table 2). Potencies against off-target sodium channel isoforms (hNa_V_1.1–1.6, hNa_V_1.8) were assessed following a similar protocol. ST-2262 was determined to be >200-fold selective over hNa_V_1.6 (IC_50_ = 17.9 μM, 95% CI 14.8–22.1), >900-fold selective over hNa_V_1.3 (IC_50_ = 65.3 μM, 62.7–68.1), and >1000-fold selective over all other Na_V_ isoforms tested. Similar IC_50_ values against hNa_V_ subtypes were obtained in an independent study using the PatchXpress automated electrophysiology platform (Suppl Table 3).

**Figure 3.**
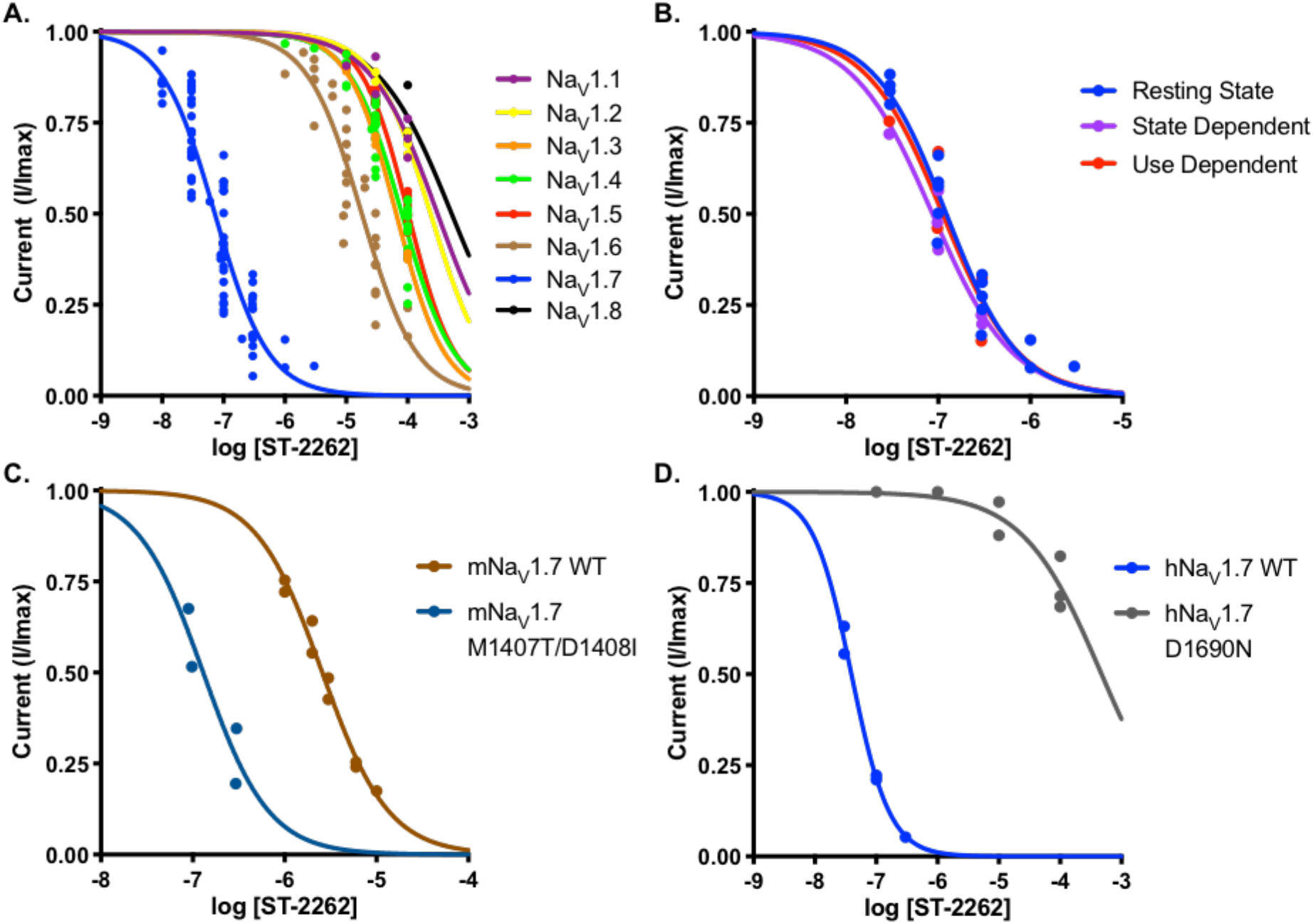
**(a)** Dose-response curves for the inhibitory effect of ST-2262 on Na_V_1.1–Na_V_1.8 stably expressed in CHO or HEK293 cells using a resting state protocol with a 10 ms pulse from rest at –110 mV to voltage at peak activation (from –20 to +10 mV). Na_V_1.X IC_50_ (in μM, mean, 95% CI). Na_V_1.1: >100; Na_V_1.2: >100; Na_V_1.3: 65.3, 62.7–68.1; Na_V_1.4: 80.7, 71.1–93.3; Na_V_1.5: >100; Na_V_1.6: 17.9, 14.8–22.1; Na_V_1.7: 0.072, 0.064–0.082; Na_V_1.8: >100. **(b)** Comparison of dose-response relationship of ST-2262 inhibition against Na_V_1.7 on different protocols. Several batches of ST-2262 were tested with different protocols. State-dependent protocol contained an 8 s conditioning step to voltage at half-inactivation, followed by a 10 ms step to voltage at full activation.(16) Use-dependent protocol contained the same step as resting-state protocol, stimulated at 30 Hz. Na_V_1.7 IC_50_ (in μM, mean, 95% CI). resting: 0.123, 0.104–0.145; state-dependent: 0.087, 0.056–0.120; use-dependent: 0.112, 0.015–0.357. **(c)** Comparison of dose-response relationship of Na_V_1.7 inhibition against WT mNa_V_1.7 and M1407T/D1408I mNa_V_1.7 on a resting state protocol. mNa_V_1.7 IC_50_ (in μM, 95% CI). WT: 2.57, 2.30–2.87; M1407T/D1408I: 0.130, 0.055–0.307. **(d)** Comparison of dose-response of ST-2262 against transiently expressed hNa_V_1.7 WT and hNa_V_1.7 D1690N. IC_50_ (in μM, mean, 95% CI). WT: 0.039, 0.032–0.047; D1690N: >100.

To investigate whether the potency of ST-2262 was dependent on the membrane holding potential (state-dependent) or frequency of stimulus (frequency-dependent), the IC_50_ against hNa_V_1.7 was measured using a protocol that favors the inactivated state of the channel and with a protocol that depolarizes the cell at high frequency. The IC_50_ of ST-2262 was not appreciably altered in the state-dependent protocol that incorporated an 8 second step to the voltage of half-inactivation (IC_50_ = 0.087 μM, 0.056–0.120), or in the high frequency protocol using a 30 Hz stimulus (IC_50_ = 0.112 μM, 0.015–0.357; Figure 3b, Suppl Table 4). These results confirm that ST-2262 is a potent and selective state- and frequency-independent inhibitor of hNa_V_1.7.

### Species variation in potency and mutagenesis

The potency of ST-2262 was assessed against a panel of species variants of Na_V_1.7 that included mouse, rat, and cynomolgus monkey channels (Suppl Table 5). Consistent with the hypothesis that Na_V_1.7 potency is affected by the presence of the 1398T/1399I sequence variation in the DIII pore loop, the IC_50_ against cynoNa_V_1.7 (0.101 μM, 0.073–0.140) was similar to human. In contrast, ST-2262 was >50x less potent against mouse (IC_50_ = 3.78 μM, 3.23-4.43) and rat Na_V_1.7 (IC_50_ = 4.95 μM, 4.17-5.87) than the human ortholog. Affinity was restored within twofold of the hNa_V_1.7 potency by introduction of domain III MD-TI mutations to mouse Na_V_1.7 (IC_50_ = 0.130 μM, 0.055–0.307; Figure 3c, Suppl Table 6).

Multiple lines of evidence suggest that ST-2262 binds to the extracellular vestibule of the sodium channel (i.e., Site 1) including: i) the structural similarity of ST-2262 to natural bis-guanidinium toxin ligands, ii) the state- and frequency-independent mode of Na_V_ inhibition that is characteristic of extracellular pore blockers, and iii) the influence of the DIII pore loop sequence on potency. To gain additional support that ST-2262 binds to the outer pore of Na_V_, we generated a point mutant of hNa_V_1.7, D1690N, at a position known to significantly destabilize binding of STX.(39) The domain IV residue D1690 forms a critical bridged hydrogen bond with the C12 hydrated ketone of STX.(39, 41) The introduction of other point mutations to Na_V_1.7 was attempted (Y362S and E916A), but these variants proved challenging to express.(39, 42) ST-2262 exhibited a >1000-fold loss in potency against hNa_V_1.7 D1690N (IC_50_ >100 μM) compared to the wild-type channel (IC_50_ = 0.039 μM, 0.032–0.047; Figure 3d, Suppl Table 7). Collectively, these results indicate that ST-2262 binds to the extracellular vestibule of Na_V_1.7, exhibits significant species variation in potency, and displays isoform selectivity in part due to molecular interactions with residues T1398 and I1399 that are unique to human and non-human primate Na_V_1.7 orthologs.(31, 32)

### ST-2262 is analgesic in a nonhuman primate model of thermal pain

Mice and humans with genetic Na_V_1.7 loss-of-function are profoundly insensitive to noxious heat.(2, 3, 43–45) To determine if acute exposure to ST-2262 can recapitulate this phenotype and in order to study the pharmacokinetic/pharmacodynamic (PK/PD) relationship of a Na_V_1.7 state-independent inhibitor, we evaluated the effect of ST-2262 in a non-human primate (NHP) model of acute thermal pain. It is not possible to study the effect of ST-2262 on acute thermal pain in rodents as this compound is >50-fold less potent against Na_V_1.7 in species that lack the T1398/I1399 sequence variant (Suppl Table 5). A NHP model of acute thermal pain was identified that uses a heat lamp to deliver a thermal stimulus to the dorsal surface of the hand of lightly anesthetized cynomolgus macaques and measures the time to withdrawal.(46) Prior to advancing ST-2262 into the NHP acute thermal pain model, a standard battery of preclinical assays was completed to evaluate ADME and pharmacokinetic properties in cynomolgus macaques (Suppl Table 7) as well as off-target activity of this inhibitor.

Male cynomolgus monkeys were anesthetized with propofol to a level in which the withdrawal reflex of the hand occurred at a consistent latency of approximately 3 seconds, a response time that was comparable to the detection of sharp pain from Aδ fibers when tested on human volunteers. The dorsal surface of the hand was exposed to a thermal stimulus that selectively activates Aδ-fiber nociceptors (Suppl Figure 4).(46, 47) The thermal stimulus was turned off at 5 seconds to prevent tissue damage; this duration produced very strong to intolerable pain in human volunteers (data not shown). Heart rate was monitored throughout the study, and presentation of the noxious thermal stimuli consistently led to a transient increase in heart rate that peaked seconds after the stimulus and then returned to baseline. Acute noxious thermal stimuli transiently increase heart rate in human subjects; the percent change in heart rate correlates with subjective pain score.(48)

ST-2262 hydrochloride administered IV dose-dependently increased the withdrawal latency to thermal stimuli with an ED_50_ of 0.13 mg/kg (Figure 4, Suppl Figure 4). At a dose of 1.25 mg/kg, all monkeys tested showed no hand withdrawal prior to the 5 second cut-off latency. Furthermore, ST-2262 dose-dependently reduced the transient heart rate increase induced by the stimuli, and the 1.25 mg/kg dose nearly abolished the effect (Figure 4b, Suppl Figure 4b). The duration of time following dosing to assess the PK/PD relationship was limited by the maximum time under propofol anesthesia permitted by our animal protocol; however, we found that 0.25 mg/kg ST-2262 resulted in ~1400 ng/ml in plasma at the 5 minute time point (n=2), which corresponds to 7x the IC_50_ against cyno Na_V_1.7 corrected for plasma protein binding (cyno PPB = 73.5%). The unbound exposure was reduced to 3.4x cyno Na_V_1.7 IC_50_ at the 30 minute time point. At a dose of 1.25 mg/kg, the total plasma concentration was ~7000 ng/ml at 5 minutes (n=2), which corresponds to an unbound exposure of 32x cynoNa_V_1.7 IC_50_, and was maintained above 15x Na_V_1.7 IC_50_ for over 100 minutes (Figure 4c). Lumbar CSF samples collected from two animals indicated that ST-2262 was peripherally restricted, with CSF:plasma ratios < 10^−3^ (n=2).

**Figure 4.**
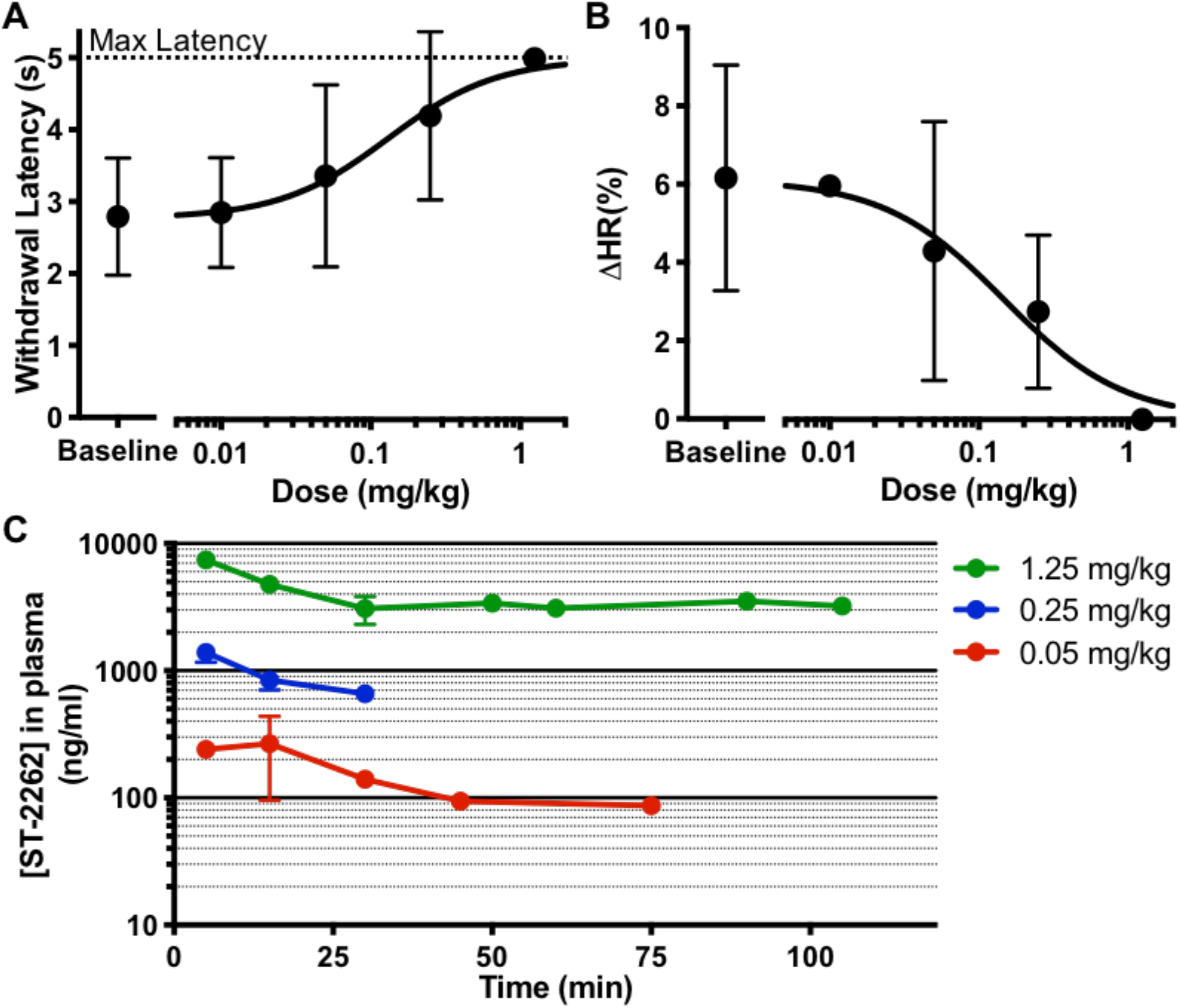
ST-2262 displays analgesic properties in a non-human primate noxious heat model. **(a)** ST-2262 induces a dose-dependent increase in the withdrawal latency to a thermal stimulus (mean ±SD; n=4). ED_50_ of effect on withdrawal latency is (in mg/kg; mean, 95% CI) 0.13, 0.055–0.28. Individual subject data shown in Suppl Figure 4. **(b)** ST-2262 dose-dependently reduces the increase in heart rate induced by the thermal stimulus (mean ±SD; n=4). ED_50_ of effect on reducing the transient heart rate increase is (in mg/kg; mean, 95% CI) 0.15, 0.061–0.32. **(c)** Plasma level concentration of ST-2262 in plasma at different doses.

By adjusting the radiant heat parameters, the NHP noxious heat model can also be used to selectively assess responses to cutaneous C-fiber nociceptor activation, which produces a burning pain in volunteers. The effect of ST-2262 on C-fiber induced hand withdrawal and heart rate change was investigated on two cynomolgus subjects.(46) As with the Aδ nociceptive response, 1.25 mg/kg ST-2262 completely abolished the C-fiber mediated hand withdrawal and heart rate change (Suppl Figure 5). Collectively, these results establish that a state-independent inhibitor of Na_V_1.7 can reduce sensitivity to noxious heat, phenotypically analogous to studies of Na_V_1.7 loss-of-function in CIP patients.(2) In addition, analysis of the PK/PD relationship of ST-2262 in this model provides insight into the level of Na_V_1.7 target occupancy that is necessary to achieve a robust PD effect.

## Discussion

The finding that humans lacking functional Na_V_1.7 exhibit an inability to experience pain raises the intriguing possibility that selective inhibitors of Na_V_1.7 may be potent analgesics.(1–3) In the present study, we describe the discovery and development of ST-2262, a selective, state- and frequency-independent inhibitor of hNa_V_1.7 that was advanced through rational modification of a natural small molecule toxin lead, STX. The properties of ST-2262 are in stark contrast to other preclinical and clinical inhibitors of Na_V_1.7, which preferentially inhibit the inactivated conformation(s) of the channel.(49). Mutagenesis experiments indicate that specific residues in the extracellular pore of Na_V_1.7, including a two-amino acid sequence variation in the domain III pore loop that is unique to primates, are required for the high potency of ST-2262 against cyno- and human Na_V_1.7.(31, 39) These findings establish the extracellular vestibule of Na_V_1.7 as a viable receptor site for the design of potent and selective channel inhibitors.

Whereas congenital insensitivity to pain in humans is the result of complete and permanent Na_V_1.7 loss-of-function, inhibition by small molecule agents is incomplete and transient. This difference raises several important questions regarding the pharmacology of Na_V_1.7: 1) is transient inhibition sufficient for analgesia, 2) what level of target engagement is required for efficacy, and 3) what anatomic compartment(s) must be accessed? Results from the present study give insights to these questions. Reduction in sensitivity to noxious thermal stimuli in NHPs occurs within minutes of IV drug administration, faster than can be explained by changes in gene expression. This observation is consistent with the view that transient inhibition of Na_V_1.7 is sufficient to produce analgesia.(44) Na_V_1.7 is present along the entire length of primary afferent nociceptive neurons from the peripheral terminals to the central terminals within the spinal dorsal horn, raising the question of whether access to the central terminals is required for analgesia.(50) ST-2262 is a polar small molecule with low membrane permeability and is unlikely to cross the blood-brain barrier to reach efficacious concentrations in the CNS. Consistent with these assumptions, analysis of CSF samples obtained during NHP experiments gave a CSF:plasma ratio of < 10^−3^. Accordingly, the reduced sensitivity to noxious stimulus of primates dosed with ST-2262 is likely the result of peripheral inhibition of Na_V_1.7.

In preclinical pain models, the majority of reported Na_V_1.7 inhibitors require high unbound plasma exposures (>20–100x the in vitro IC_50_ against Na_V_1.7) to achieve a pharmacodynamic effect.(8) Examples include a cystine knot toxin that binds to VSD II and select sulfonamide inhibitors that bind to VSD IV.(44, 51–53). For this latter class of inhibitors, determining the level of Na_V_1.7 target occupancy reached at these exposures is difficult given the variation in potency (i.e., IC_50_ values determined by electrophysiology) against resting and inactivated conformational states of the channel.(49) In contrast, ST-2262 displays the same IC_50_ against both the rest and inactivated states of Na_V_1.7, and this value is not influenced by the frequency of channel opening. Consequently, with ST-2262, correlating the level of Na_V_1.7 target occupancy *in vivo* with the measured unbound plasma concentration is straightforward, as is the percent Na_V_1.7 inhibition with measured pharmacodynamic effects.

In the present study, the effect of ST-2262 on withdrawal latency to noxious heat was measured in NHPs at doses of 0.05, 0.25 and 1.25 mg/kg IV. These doses give unbound plasma concentrations of ST-2262 of 0.7x, 3.4x and 16x the IC_50_ against cynoNa_V_1.7 at a time point 30 minutes after drug administration. Assuming a Hill slope of 1, which is consistent with the doseresponse relationship observed for ST-2262 in whole cell recordings against cyno and human Na_V_1.7, these unbound exposures correspond to 41%, 78% and 94% inhibition of Na_V_1.7, respectively. Our findings thus support the view that a relatively high but not complete level of Na_V_1.7 inhibition (~70%) is required for robust analgesia in preclinical pain models.(8)

## Conclusion

Despite recent clinical setbacks, Na_V_1.7 remains a compelling target for the development of non-opioid analgesics based on evidence from human genetics and rodent KO studies.(2, 3, 43, 44) A major challenge in the pursuit of safe and effective Na_V_1.7 inhibitors has been the identification of small molecule inhibitors with sufficient selectivity over off-target Na_V_ isoforms to achieve a suitable margin of safety. Prior efforts to develop Na_V_1.7-selective small molecules have largely focused on a class of aryl and acyl sulfonamides that bind to VSD IV and preferentially inhibit an inactivated conformational state of the channel.(7) In the present study, we disclose ST-2262, a highly selective synthetic analogue of natural bis-guanidinium toxins that binds to the extracellular vestibule of the channel (Site 1) and occludes ion passage. When administered systemically, ST-2262 recapitulates a distinguishing characteristic of individuals genotyped with Na_V_1.7 loss-of-function – reduced sensitivity to noxious heat. These findings validate the extracellular mouth of the sodium channel as a tractable binding site for selective ligand design and provide insight into the distribution and target occupancy requirements for analgesia mediated by inhibition of Na_V_1.7.

## References

1. S. D. Dib-Hajj, Y. Yang, J. A. Black, S. G. Waxman, The Na(V)1.7 sodium channel: from molecule to man. Nat. Rev. Neurosci. 14, 49–62 (2013).

2. J. J. Cox, et al., An SCN9A channelopathy causes congenital inability to experience pain. Nature 444, 894–898 (2006).

3. Y. Goldberg, et al., Loss-of-function mutations in the Nav1.7 gene underlie congenital indifference to pain in multiple human populations. Clinical Genetics 71, 311–319 (2007).

4. Y. Yang, et al., Mutations in SCN9A, encoding a sodium channel alpha subunit, in patients with primary erythermalgia. J. Med. Genet. 41, 171–174 (2004).

5. C. G. Faber, et al., Gain of function Nav1.7 mutations in idiopathic small fiber neuropathy. Ann. Neurol. 71, 26–39 (2012).

6. J. Payandeh, D. H. Hackos, “Selective Ligands and Drug Discovery Targeting the Voltage-Gated Sodium Channel Nav1.7 “ in Voltage-Gated Sodium Channels: Structure, Function and Channelopathies, M. Chahine, Ed. (Springer International Publishing, 2018), pp. 271–306.

7. S. J. McKerrall, D. P. Sutherlin, Nav1.7 inhibitors for the treatment of chronic pain. Bioorganic & Medicinal Chemistry Letters 28, 3141–3149 (2018).

8. J. V. Mulcahy, et al., Challenges and Opportunities for Therapeutics Targeting the Voltage-Gated Sodium Channel Isoform NaV1.7. J. Med. Chem. (2019) https:/doi.org/10.1021/acs.jmedchem.8b01906.

9. C. A. Ahern, J. Payandeh, F. Bosmans, B. Chanda, The hitchhiker’s guide to the voltage-gated sodium channel galaxy. J. Gen. Physiol. 147, 1–24 (2016).

10. M. Stevens, S. Peigneur, J. Tytgat, Neurotoxins and their binding areas on voltage-gated sodium channels. Front Pharmacol 2, 71 (2011).

11. M. R. Israel, B. Tay, J. R. Deuis, I. Vetter, “Sodium Channels and Venom Peptide Pharmacology “ in Advances in Pharmacology, (Elsevier, 2017), pp. 67–116.

12. F. Bosmans, K. J. Swartz, Targeting voltage sensors in sodium channels with spider toxins. Trends in Pharmacological Sciences 31, 175–182 (2010).

13. X. Wang, et al., Inhibitors of Ion Channels, Patent PCT/US2006/042882, 2006.

14. A. Fulp, B. Marron, M. Suto J., X. Wang, Inhibitors of Voltage-Gated Sodium Channels, Patent PCT/US2006/031390, 2006.

15. A. Kawatkar S., et al., Bicyclic Deriatives as Modulators of Ion Channels, Patent PCT/US2006/017699, 2006.

16. K. McCormack, et al., Voltage sensor interaction site for selective small molecule inhibitors of voltage-gated sodium channels. Proceedings of the National Academy of Sciences 110, E2724–E2732 (2013).

17. M.-M. Zhang, et al., Cooccupancy of the outer vestibule of voltage-gated sodium channels by micro-conotoxin KIIIA and saxitoxin or tetrodotoxin. J. Neurophysiol. 104, 88–97 (2010).

18. H. Shen, et al., Structural basis for the modulation of voltage-gated sodium channels by animal toxins. Science 362, eaau2596 (2018).

19. S. Ahuja, et al., Structural basis of Nav1.7 inhibition by an isoform-selective small-molecule antagonist. Science 350, aac5464 (2015).

20. A. J. Alexandrou, et al., Subtype-Selective Small Molecule Inhibitors Reveal a Fundamental Role for Nav1.7 in Nociceptor Electrogenesis, Axonal Conduction and Presynaptic Release. PLoS ONE 11, e0152405 (2016).

21. M. Varney, Roche: At the Forefront of R&D Innovation and Breakthrough Treatments (2016).

22. W. A. Schmalhofer, et al., ProTx-II, a selective inhibitor of NaV1.7 sodium channels, blocks action potential propagation in nociceptors. Mol. Pharmacol. 74, 1476–1484 (2008).

23. Y. Xiao, et al., Tarantula hnwentoxin-IV inhibits neuronal sodium channels by binding to receptor site 4 and trapping the domain ii voltage sensor in the closed configuration. J. Biol. Chem. 283, 27300–27313 (2008).

24. J. R. Deuis, et al., Pharmacological characterisation of the highly NaV1.7 selective spider venom peptide Pn3a. Sci Rep 7, 40883 (2017).

25. B. D. Moyer, et al., Pharmacological characterization of potent and selective NaV1.7 inhibitors engineered from Chilobrachys jingzhao tarantula venom peptide JzTx-V. PLoS ONE 13, e0196791 (2018).

26. M. Flinspach, et al., Insensitivity to pain induced by a potent selective closed-state Nav1.7 inhibitor. Sci Rep 7, 39662 (2017).

27. S. J. McKerrall, et al., Structure-and Ligand-Based Discovery of Chromane Arylsulfonamide Nav1.7 Inhibitors for the Treatment of Chronic Pain. J. Med. Chem. 62, 4091–4109 (2019).

28. R. F. Graceffa, et al., Sulfonamides as Selective NaV1.7 Inhibitors: Optimizing Potency, Pharmacokinetics, and Metabolic Properties to Obtain Atropisomeric Quinolinone (AM-0466) that Affords Robust in Vivo Activity. J. Med. Chem. 60, 5990–6017 (2017).

29. A. McDonnell, et al., Efficacy of the Nav1.7 blocker PF-05089771 in a randomised, placebo-controlled, double-blind clinical study in subjects with painful diabetic peripheral neuropathy. Pain 159, 1465–1476 (2018).

30. M. E. Rothenberg, et al., Safety, Tolerability, and Pharmacokinetics of GDC-0276, a Novel NaV1.7 Inhibitor, in a First-in-Human, Single-and Multiple-Dose Study in Healthy Volunteers. Clin Drug Investig (2019) https:/doi.org/10.1007/s40261-019-00807-3.

31. J. R. Walker, et al., Marked difference in saxitoxin and tetrodotoxin affinity for the human nociceptive voltage gated sodium channel (Nav1.7) [corrected]. Proc. Natl. Acad. Sci. U.S.A. 109, 18102–18107 (2012).

32. R. Thomas-Tran, J. Du Bois, Mutant cycle analysis with modified saxitoxins reveals specific interactions critical to attaining high-affinity inhibition of hNaV1.7. Proc. Natl. Acad. Sci. U.S.A. 113, 5856–5861 (2016).

33. T. Tsukamoto, et al., Differential binding of tetrodotoxin and its derivatives to voltage-sensitive sodium channel subtypes (Nav 1.1 to Nav 1.7). Br. J. Pharmacol. 174, 3881–3892 (2017).

34. J. J. Fleming, M. D. McReynolds, J. Du Bois, (+)-saxitoxin: a first and second generation stereoselective synthesis. J. Am. Chem. Soc. 129, 9964–9975 (2007).

35. J. V. Mulcahy, J. R. Walker, J. E. Merit, A. Whitehead, J. Du Bois, Synthesis of the Paralytic Shellfish Poisons (+)-Gonyautoxin 2, (+)-Gonyautoxin 3, and (+)-11,11-Dihydroxysaxitoxin. J. Am. Chem. Soc. 138, 5994–6001 (2016).

36. B. M. Andresen, J. Du Bois, De novo synthesis of modified saxitoxins for sodium ion channel study. J. Am. Chem. Soc. 131, 12524–12525 (2009).

37. J. R. Walker, J. E. Merit, R. Thomas-Tran, D. T. Y. Tang, J. Du Bois, Divergent Synthesis of Natural Derivatives of (+)-Saxitoxin Including 11-Saxitoxinethanoic Acid. Angew. Chem. Int. Ed. Engl. 58, 1689–1693 (2019).

38. G. M. Lipkind, H. A. Fozzard, A structural model of the tetrodotoxin and saxitoxin binding site of the Na+ channel. Biophys. J. 66, 1–13 (1994).

39. J. L. Penzotti, H. A. Fozzard, G. M. Lipkind, S. C. Dudley, Differences in saxitoxin and tetrodotoxin binding revealed by mutagenesis of the Na+ channel outer vestibule. Biophys. J. 75, 2647–2657 (1998).

40. D. B. Tikhonov, B. S. Zhorov, Modeling P-loops domain of sodium channel: homology with potassium channels and interaction with ligands. Biophys. J. 88, 184–197 (2005).

41. H. Shen, D. Liu, K. Wu, J. Lei, N. Yan, Structures of human Nav 1.7 channel in complex with auxiliary subunits and animal toxins. Science 363, 1303–1308 (2019).

42. J. Satin, et al., A Mutant of TTX-Resistant Cardiac Sodium Channels with TTX-Sensitive Properties. Science 256, 1202–1205 (1992).

43. J. Gingras, et al., Global Nav1.7 knockout mice recapitulate the phenotype of human congenital indifference to pain. PLoS ONE 9, e105895 (2014).

44. S. D. Shields, et al., Insensitivity to Pain upon Adult-Onset Deletion of Nav1.7 or Its Blockade with Selective Inhibitors. J. Neurosci. 38, 10180–10201 (2018).

45. M. S. Minett, et al., Distinct Nav1.7-dependent pain sensations require different sets of sensory and sympathetic neurons. Nat Commun 3, 791 (2012).

46. D. C. Yeomans, et al., Recombinant herpes vector-mediated analgesia in a primate model of hyperalgesia. Molecular Therapy 13, 589–597 (2006).

47. D. C. Yeomans, V. Pirec, H. K. Proudfit, Nociceptive responses to high and low rates of noxious cutaneous heating are mediated by different nociceptors in the rat: behavioral evidence. Pain 68, 133–140 (1996).

48. M. L. Loggia, M. Juneau, C. M. Bushnell, Autonomic responses to heat pain: Heart rate, skin conductance, and their relation to verbal ratings and stimulus intensity: Pain 152, 592–598 (2011).

49. J. W. Theile, M. D. Fuller, M. L. Chapman, The Selective Nav1.7 Inhibitor, PF-05089771, Interacts Equivalently with Fast and Slow Inactivated Nav1.7 Channels. Mol. Pharmacol. 90, 540–548 (2016).

50. J. A. Black, N. Frézel, S. D. Dib-Hajj, S. G. Waxman, Expression of Nav1.7 in DRG neurons extends from peripheral terminals in the skin to central preterminal branches and terminals in the dorsal horn. Mol Pain 8, 82 (2012).

51. B. Wu, et al., Discovery of Tarantula Venom-Derived NaV1.7-Inhibitory JzTx-V Peptide 5-Br-Trp24 Analogue AM-6120 with Systemic Block of Histamine-Induced Pruritis. J. Med. Chem. 61, 9500–9512 (2018).

52. J. E. Pero, et al., Benzoxazolinone aryl sulfonamides as potent, selective Na v 1.7 inhibitors with in vivo efficacy in a preclinical pain model. Bioorganic & Medicinal Chemistry Letters 27, 2683–2688 (2017).

53. T. J. Kornecook, et al., Pharmacologic Characterization of AMG8379, a Potent and Selective Small Molecule Sulfonamide Antagonist of the Voltage-Gated Sodium Channel NaV1.7. J. Pharmacol. Exp. Ther. 362, 146–160 (2017).

