## Supporting Information for "Discovery of a Selective, State-Independent Inhibitor of Na_V_1.7 by Modification of Guanidinium Toxins"

**Supplementary Materials:**

Materials and Methods

**Guanidinium Toxin Synthesis.** dcSTX, ST-282, and ST-2262 were prepared from L-serine methyl ester by modification of a previously reported synthetic route.(35, 37) All reagents were obtained commercially unless otherwise noted. Final compounds were purified by reversed-phase HPLC using a Shimazdu LC-20AP purification system and Bonna-Agela Durashell C18 column, and characterized by NMR spectroscopy using a Bruker Ultrashield Plus 400 MHz spectrometer. Characterization information can be found in *SI Materials and Methods*.

**Mutagenesis.** Primers were synthesized by Eton Biosciences (h$Na_V1.7$ primers) or IDT (m$Na_V1.7$ M1407T/D1408I). Primer sequences can be found in SI Materials and Methods. Mutants were prepared using the QuikChange Lightning Site-Directed Mutagenesis Kit (Agilent Technologies) according to the manufacturer's protocols. Mutations were confirmed by DNA sequencing (Eton Biosciences or Genewiz) of the entire resulting plasmids.

Forward primer sequences (5′ to 3′) used for site-directed mutagenesis of h$Na_V1.7$:

| | |
|---|---|
| **E364Q** | GACCCAAGATTACTGGCAAAACCTTTACCAACAGACG |
| **E916A** | GTGCTGTGTGGAGCGTGGATAGAGACCATGTGG |
| **T1398M** | GCAACTTTTAAGGGATGGATGATTATTATGTATGCAGCAGTGG |
| **T1398M/I1399D** | GCAACTTTTAAGGGATGGATGGATATTATGTATGCAGCAGTGG |
| **D1690N** | CCTCTGCTGGCTGGAATGGATTGCTAGCACC |

Forward primer sequences (5' to 3') used for site-directed mutagenesis of m$Na_V1.7$ (Transomic, clone BC172147 in pTCN vector):

**M1407T/D1408I**
GTTGCAACGTTCAAGGGCTGGACGATTATTATGTATGCAGCAGTTGAC

**Reverse primers are the reverse complement of the corresponding forward primer.**

**Cell Culture and Manual Patch Clamp Electrophysiology.** Whole cell recordings were carried out on human embryonic kidney 293 (HEK) or Chinese Hamster Ovary (CHO) cells that stably expressed one $Na_V1.x$ isoform. The following cell lines were utilized: $Na_V1.1$ – CHO, division arrested, purchased from Charles River Laboratories (CRL, ChanTest, Cleveland, OH), Catalog #CT4178; $Na_V1.2$ – CHO, division arrested, CRL, Catalog #CT4010; $Na_V1.3$ – CHO, division arrested, CRL, Catalog #CT4157; $Na_V1.4$ – HEK, shared by academic collaborator, see Mitrovic et al, 1996; $Na_V1.5$ – HEK, SB Drug Discovery (Glasgow, UK), Catalog #SB-HEK-h$Na_V1.5$; $Na_V1.6$ - HEK, SB Drug Discovery. Catalog #SB-HEK-h$Na_V1.6$; $Na_V1.7$ - HEK, SB Drug Discovery. Catalog #SB-HEK-h$Na_V1.7$; $Na_V1.8$ – CHO, division arrested, CRL, Catalog #CT4011. Stably transfected competent cells were maintained in Dulbecco's Modified Eagle's

Medium (DMEM; Thermo Fisher, Waltham MA) supplemented with 10% fetal bovine serum (Thermo Fisher), 100 U/mL penicillin G sodium, 100 μg/mL streptomycin sulfate (VW, Radnor, PA), 500 μg/mL G418 (Thermo Fisher), and blasticidin S hydrochloride (for cell lines from SB Ion Channels; VWR). Division arrested cells were maintained for 2–3 days in Ham's Nutrient Mixture F-12 (VWR) media supplemented with 10% fetal bovine serum, 100 U/mL penicillin G sodium, 100 μg/mL streptomycin sulfate.

For recordings with mouse $Na_V1.7$ and $Na_V1.7$ mutant channels, HEK or CHO cells were transiently transfected with plasmids encoding the corresponding genes. The following transgenes were utilized: Scn9a *Mus musculus* – TransOMIC (Huntsville, AL), Clone #BC172147; pcDNA3.1(+)IRES GFP? – Addgene (Watertown, MA), Catalog #51406. CHO or HEK cells were plated into a 96-well plate at 80–90% confluency. The cells were rinsed in phosphate-buffered saline (PBS) followed by the addition of opti-MEM media (Thermo Fisher) and the transfection mix with reagents from the Lipofectamine 3000 kit (Thermo Fisher). Transfection mix contained the plasmid encoding the gene of interest along with the plasmid encoding green fluorescent protein (GFP). Cells were incubated for 3 days to allow the heterologous expression. GFP expression was used to identify transfected cells on the day of the experiment.

On the day of testing, cells were rinsed with PBS, lifted from the plate with Detachin (VWR), resuspended in fresh media, and 30 μl were plated onto 5 mm glass cover slips (Bellco Glass, Vineland, NJ). After cells adherence, the cover slip was transferred to the testing chamber and continuously superfused with extracellular solution, which contained (in mM): NaCl (135), KCl (4.5) $CaCl_2$ (2), $MgCl_2$ (1), HEPES (10); pH 7.4. Cells were patch clamped with borosilicate glass pipettes pulled to a tip diameter yielding a resistance of 1.0–2.0 MΩ, and filled with an internal solution that contained (in mM): CsF (125), NaCl (10), EGTA (10), HEPES (10), pH 7.2. ST-2262 was dissolved in DMSO in a 10 mM stock and then subsequently was serially diluted in the extracellular solution. Alone, the maximum DMSO concentration of 1% (in the 100 uM ST-2262 test solution) had no effect on sodium currents (data not shown).

Channel currents were measured using whole-cell patch-clamp electrophysiology with a HEKA EPC 9 amplifier with built-in ITC-16 interface (HEKA Elektronik Dr. Schulze GmbH, Germany). The output of the EPC 9 patch-clamp amplifier was filtered with a built-in low-pass, fourpole Bessel filter with a cutoff frequency of 10 kHz and was sampled at 20–50 kHz. Pulse stimulation and data acquisition were controlled with the Pulse software (HEKA Elektronik Dr. Schulze GmbH, Germany v8.40). All measurements were performed at room temperature (20–22°C). Recordings began at least 5 min after establishing the whole-cell and voltage-clamp configuration. Cells with access resistance < 5 MΩ were used for the study. Series resistance compensation circuit was turned on and set at 80% and 100 μsec.

Cells were held at –90–110 mV, and the voltage dependence of activation ($V_{act}$) was determined with a series of voltage steps. The voltage step that elicited the maximal current (–20 to +10mV depending on the isoform) was subsequently used to evoke a channel current from its resting state. Currents were evoked every 3 seconds with a depolarizing step to the voltage of maximal activation for 10 ms. For determining use-dependent block, the voltage step occurred at 30 Hz. The inactivated state-dependent (SD) inhibition was determined as previously described.(16) Briefly, the voltage of half-inactivation was determined by depolarizing the cell to 0 mV and increasing the resting potential by +10 mV in each successive step, from –110 to 0 mV. For state-dependent voltage steps, the cell was stepped to the voltage of half inactivation for 8 seconds. Then the cell was brought back to the resting potential for 20 ms and then stepped to the voltage of maximal activation for 20 ms.

Once the baseline evoked current was stable in amplitude, ST-2262 was washed onto the cell. Following stable inhibition, a half-log higher concentration was perfused onto the cell. Between 1 and 5 concentrations were washed onto each cell, with 100 μM being the maximum concentration tested. Drug was then washed out until the cell returned to a stable current. Saxitoxin (Millpore Sigma, Burlington, MA) was used as a positive control for all $Na_V1.x$ isoforms, and the concentration near the $IC_{50}$ for each isoform was tested either prior to or following ST-2262 perfusion or washout.

Recorded peak current data were baseline-normalized. The steady state before and after each test article application were used to calculate the percentage of current inhibited at each concentration. Percent inhibition as a function of compound concentration was pooled from all recorded cells and the resulting data set was fitted with a four-parameter logistic curve using least squares regression in GraphPad Prism, version 7 (San Diego, CA) to produce a single $IC_{50}$ curve and asymmetrical (likelihood) 95% confidence interval.

**PatchXpress 7000A Electrophysiology.** Automated patch clamp electrophysiology experiments were conducted as previously described (Wu et al, 2018, J Med Chem). Human $Na_V1.2$ and $Na_V1.3$ CHO, and $Na_V1.1$, $Na_V1.4$, $Na_V1.5$, $Na_V1.6$, and $Na_V1.7$ HEK stable cell lines were purchased from Eurofins Pharma Discovery Services (St. Charles, MO). All cell lines were validated by RT-PCR to express the indicated $Na_V$ isoform and mycoplasma free. Adherent cells were isolated from tissue culture flasks using 1:10 diluted 0.25% trypsin–EDTA treatment for 2–3 min and then were incubated in complete culture medium containing 10% fetal bovine serum for at least 15 min prior to resuspension in external solution consisting of (in mM) NaCl (70), KCl (5), d-mannitol (140), HEPES (10), $CaCl_2$ (2), $MgCl_2$ (1), and glucose (10), pH 7.4, with NaOH. Internal solution consisted of (in mM) CsCl (62.5), CsF (75), HEPES (10), EGTA (5), and $MgCl_2$ (2.5), pH 7.25, with CsOH. For Nav1.1, Nav1.2 and Nav1.3 cell lines, 140 mM NaCl was used in the external solution, with removal of mannitol, to increase macroscopic current sizes. Cells were voltage clamped using the whole-cell patch clamp configuration at room temperature (~22 °C) at a holding potential of −120 mV with test potentials to –10 mV (with the exception of a –20 mV test potential for Nav1.5). Test compounds were added, and $Na_V$ currents were monitored at 0.1 Hz at the appropriate test potential. Cells were used for additional compound testing if currents recovered to >80% of starting values following compound washout. Three to six different concentrations of test compound were applied to each channel tested. Percent inhibition as a function of compound concentration was calculated and the results from at least 9 compound applications were pooled and the resulting data set was fit with a four-parameter logistic curve using least squares regression in GraphPad Prism, version 7 (San Diego, CA) to produce a single $IC_{50}$ curve and asymmetrical (likelihood) 95% confidence interval.

**Plasma Protein Binding.** Rapid equilibrium dialysis (RED) device inserts along with a Teflon base plate (Pierce, Rockford, IL) were used for binding studies. Human and cynomolgus monkey plasma were obtained from BioIVT. The pH of the plasma was adjusted to 7.4 prior to the experiment.

DMSO stock (1 mM) of ST-2262 was spiked into plasma to make a final concentration of 2 μM. The spiked plasma solutions were placed into the sample chamber while the PBS buffer, pH 7.4, was placed into the adjacent buffer chamber. The plate was sealed with a self-adhesive lid and incubated at 37ºC on an orbital shaker (250 rpm) for 4 hours. After 4 hours, aliquots were taken from both the sample and buffer chamber. An equal volume of buffer was added to the plasma samples and plasma was added to the buffer samples. The resulting samples were analyzed by LC-MS/MS.

**Animal Studies**

Pharmacokinetics in cynomolgus monkey. A PK study was conducted at Wuxi Apptec (Suzhou, China). The study was conducted in accordance with the Wuxi IACUC guidelines, which are in compliance with the Animal Welfare Act, and the Guide for the Care and Use of Laboratory Animals. Three male cynomolgus monkeys over 2 years old received an IV administration of ST-2262. Blood samples were collected into collection tubes coated in $K_2$-EDTA at 0.083, 0.25, 0.5, 1, 2, 4, 8 and 24 hours post-dose. Samples were centrifuged at 3,000 x g for 10 minutes at 2 to 8° C, and then at least 0.15 ml plasma was removed for bio-analysis.

A 40 µl aliquot of the plasma sample was protein precipitated with 200 µl of the internal standard in acetonitrile (ACN). The mixture was vortexed and centrifuged at 13000 rpm for 10 minutes. Then, 2 µl supernatant was injected for LC-MS/MS analysis. Samples were analyzed on a Sciex 6500+ triple quadrupole mass spectrometer using a Turbo Spray IonDrive electrospray ion source equipped with a Waters Acquity UPLC system. Chromatographic separation was done on a ACE 3 AQ, 2.1 X 100 mm, 3 µm particle size column employing a gradient consisting of mobile phase A (0.025% formic acid and 1 mM ammonium acetate in water/ACN (v:v 95:5)) and mobile phase B (0.025% formic acid and 1 mM ammonium acetate in ACN/water (v:v 95:5)) starting at 0% B initially to 0.8 minutes followed by 30% B from 0.8 to 1.6 minutes and then 95% B from 1.6 to 2.71 minutes. Plasma concentration values (ng/ml) were imported into Phoenix WinNonlin 6.3 (Certara, Princeton, NJ) and pharmacokinetic parameters were derived using a noncompartmental model.

Non-human primate (NHP) noxious heat model. All experimental procedures were approved by Montana State University's IACUC in accordance with the National Institutes of Health's Guide for the Care and Use of Laboratory Animals, and conducted at an AAALAC-accredited facility. Experiments were conducted in accordance with the recommendations of the International Association for the Study of Pain, and all efforts were taken to minimize pain and distress of experimental animals.

The study was conducted as previously described.(46) Briefly, withdrawal responses of four male cynomolgus monkeys (*Macaca fascicularis*), aged 4-7 years, were assessed using the noxious heat model. Subjects were sedated with ketamine/dexmetatomidine (IM; 2.5 mg/kg ketamine (Zoetis, Parsippany, NJ), 0.04mg/kg dexmetatomidine (Zoetis)) and then up to two catheters were placed into the saphenous veins. One IV line was used to infuse propofol (2 mg/ml in saline; Zoetis); the other catheter was used for experimental drug infusions and blood draws. Vital signs including heart rate, ECG, pO2, temperature and respiratory rate were continuously monitored throughout the experiment using an ICU monitor.

Following reversal of dexmetatomidine with atipamezole (IM, 0.4 mg/kg; Zoetis), propofol flow was initiated. The dorsal side of each hand was then shaved and cleaned and the propofol flow rate was lowered in order to produce a light anesthetic plane, defined by the presence of repeatable withdrawal reflexes to noxious thermal stimuli, applied to the dorsal surface of the hand at a latency consistent with the those that previously had demonstrated to induce pain in human volunteers. Thermal stimulation was provided by a focused radiant heat source at producing skin temperature increase rates of either 6.5 or 0.9 ºC/s to either elicit withdrawals induced by A-delta or C nociceptive fiber activation, respectively.(40) The maximum (cut off) duration for the A-delta stimulus was 5 seconds; the maximum duration for C fiber stimulation was set at 20 seconds, both to prevent tissue damage. In addition to hand withdrawal, thermal stimuli reliably and transiently increased heart rate, assessed by recording the heart rate just prior to the stimulus and the maximum heart rate within 30 seconds post-stimulus.

Once withdrawal responses was stable, animals were dosed IV with ST-2262. Withdrawal testing began 10 minutes after the drug was administered. Escalating doses were given to each subject, and each successive dose was about 30 minutes apart. The experimenter was not blinded to dose.

To understand the relationship between drug concentration in plasma and paw withdrawal latency change, blood was collected into tubes coated in $K_2$-EDTA at various time points. Samples were centrifuged at 3000 x g for 10 minutes at 2° to 8° C, and then at least 0.15 ml plasma was removed for analysis of ST-2262 plasma levels. Plasma was analyzed for using a Shimadzu LC30 AD HPLC and AB Sciex API 5000 mass spectrometer, and reported as ng/mL.

Supplementary Figures and Tables

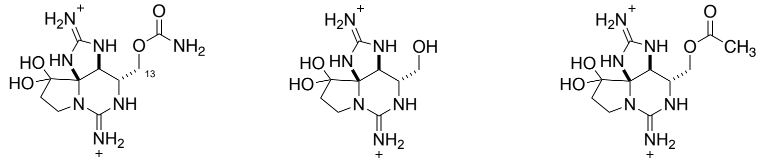


| STX | dcSTX | STX-OAc |
|---|---|---|
| $hNa_V1.7$ $IC_{50}$ 702 ± 53 nM | $hNa_V1.7$ $IC_{50}$ 186 ± 19 nM | $hNa_V1.7$ $IC_{50}$ 19 ± 1.7 nM |
| $rNa_V1.4$ $IC_{50}$ 2.9 ± 0.1 nM | $rNa_V1.4$ $IC_{50}$ 35 ± 1.4 nM | $rNa_V1.4$ $IC_{50}$ 83 ± 3.9 nM |

**Supplemental Figure 1: Differential potency of C13-modified saxitoxins.** Substitution of STX at C13 increases potency against $hNa_V1.7$ and diminishes potency against $rNa_V1.4$.(31, 32)

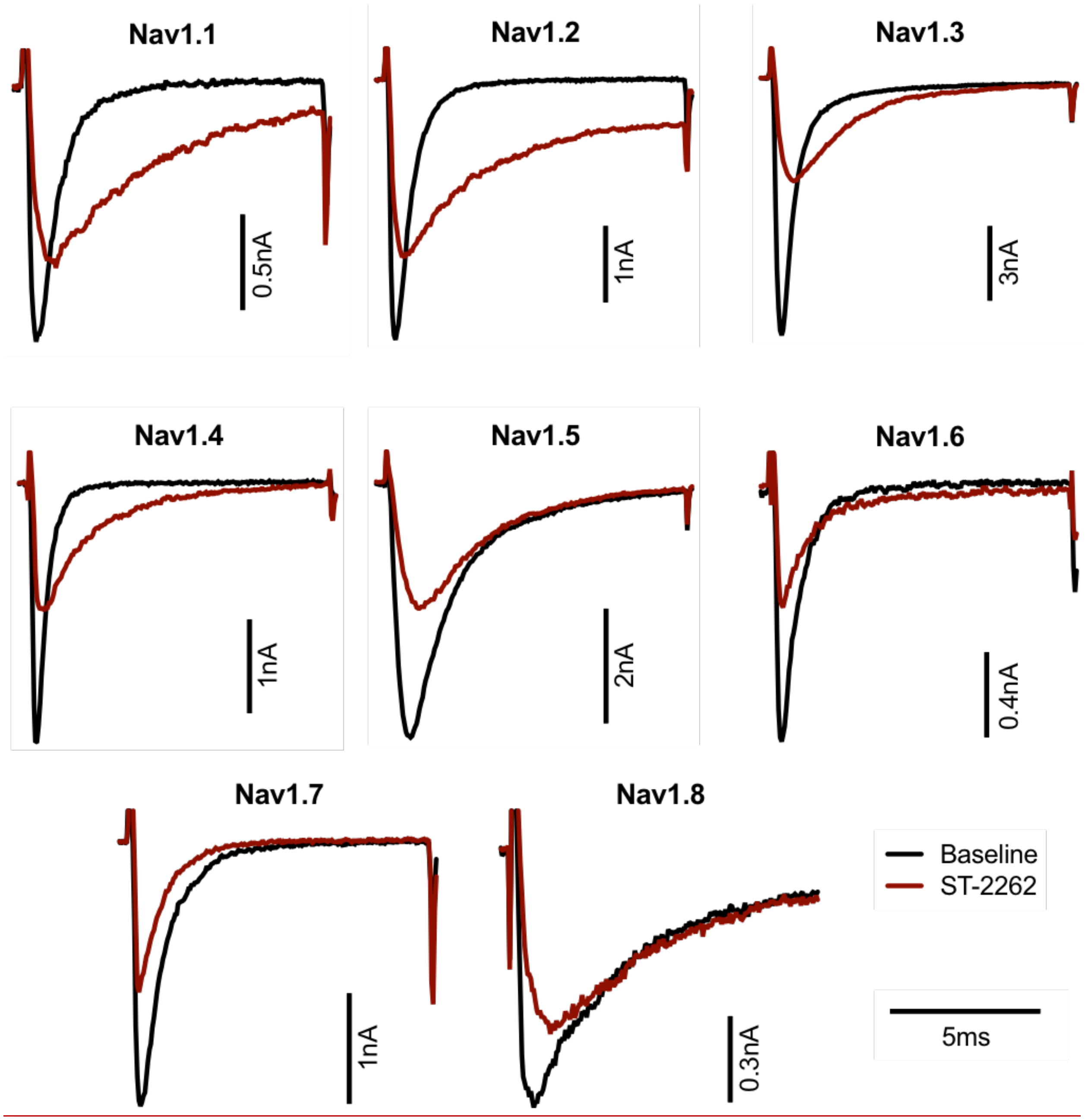


**Supplemental Figure 2: ST-2622 inhibition of Nav1.X isoforms.** Representative waveforms of $Na_V$1.X at baseline (black) and following inhibition with ST-2262 (red). Concentration of ST-2262 for the representative trace were chosen to be near $IC_{50}$. [ST-2262] is as follows. Nav1.1–Nav1.5, Nav1.8: 100 µM; Nav1.6: 30 µM; Nav1.7: 100 nM.

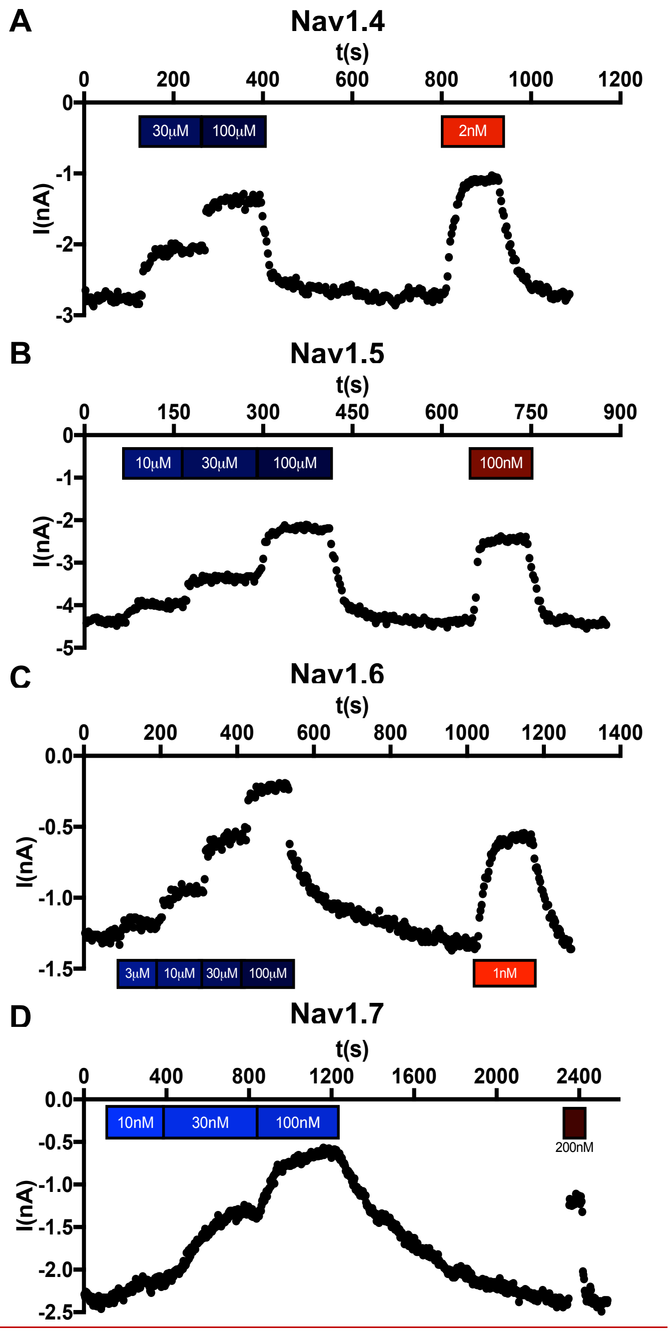


**Supplemental Figure 3: Representative time course of patch clamp recording with ST-2262 and STX inhibition.** Na$_V$1.4, Na$_V$1.5, Na$_V$1.6, and Na$_V$1.7. inhibition by ST-2262 (blue) and STX (red) bot hat various doses. STX concentration was chosen to be near $IC_{50}$ for each isoform, and was perfused onto cells following full ST-2262 washout for quality control.

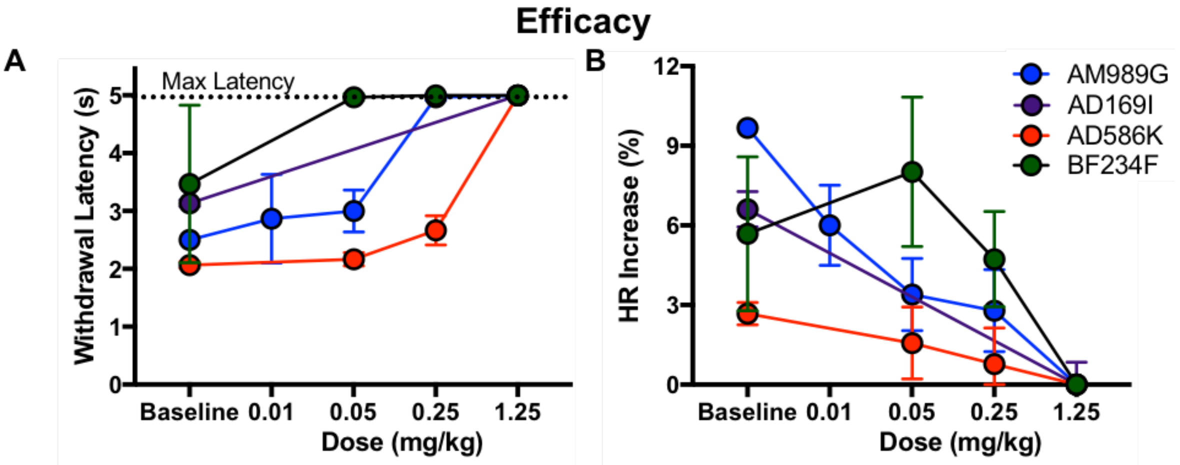


**Supplemental Figure 4: Individual subject data for the cyno efficacy study, presented as group data in Figure 4.** The efficacy endpoints were withdrawal latency **(a)** and heart rate change **(b).**

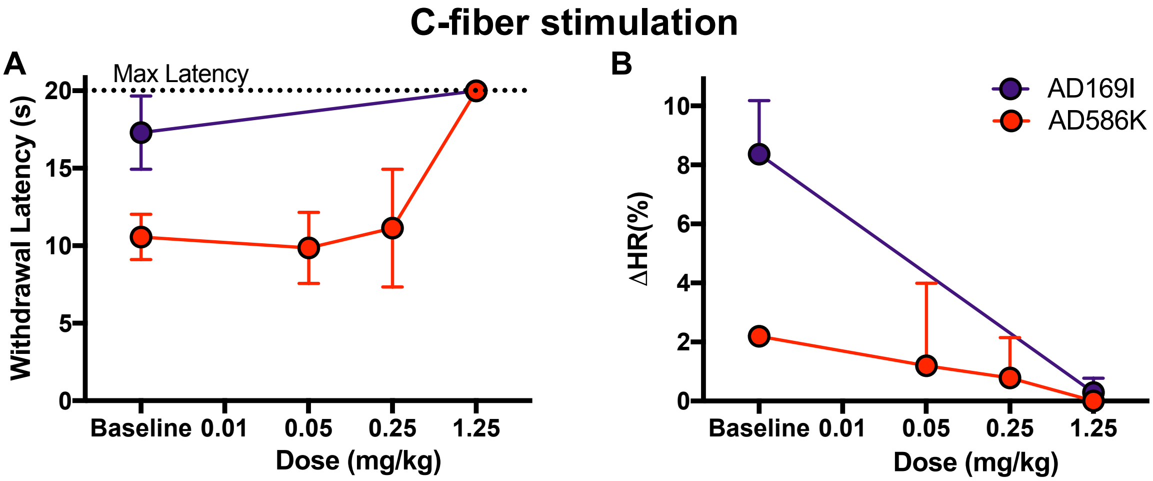


**Supplemental Figure 5: Effect of ST-2262 on withdrawal latency and heart rate change induced by C-fiber stimulation**. A lower heating rate thermal stimulus was presented for a maximum of 20 seconds, which selectively activates C fibers.(46) In two subjects, the C-fiber-induced hand withdrawal response was replicable for testing. The efficacy endpoints measured were withdrawal latency **(a)** and heart rate change **(b).** At each dose, the efficacy endpoints were repeated 3 times (presented as mean ± SD).

**Supplemental Table 1:** Sequence alignment of S5–S6 pore-forming loops from $Na_V$1.7 compared across species and alignment from human $Na_V$1.X isoforms. Key MD–TI variation in domain III is underlined.

| | Domain I | Domain II | Domain III | Domain IV | NCBI Accession |
|---|---|---|---|---|---|
| $Na_V$1.7 species variants | | | | | |
| h$Na_V$1.7 | RLMTQDYWEN | RVLCGEWIET | VATFKGWTII | ITTSAGWDGL | NP_002968 |
| cyno$Na_V$1.7 | RLMTQDYWEN | RVLCGEWIET | VATFKGWTII | ITTSAGWDGL | XP_015287803 |
| m$Na_V$1.7 | RLMTQDYWEN | RVLCGEWIET | VATFKGWMDI | ITTSAGWDGL | NP_001277603 |
| r$Na_V$1.7 | RLMTQDYWEN | RVLCGEWIET | VATFKGWMDI | ITTSAGWDGL | NP_579823 |
| dog$Na_V$1.7 | RLMTQDYWEN | RVLCGEWIET | VATFKGWMDI | ITTSAGWDGL | XP_022270547 |
| human $Na_V$1.X isoforms | | | | | |
| h$Na_V$1.1 | RLMTQDFWEN | RVLCGEWIET | VATFKGWMDI | ITTSAGWDGL | NP_001159435 |
| h$Na_V$1.2 | RLMTQDFWEN | RVLCGEWIET | VATFKGWMDI | ITTSAGWDGL | NP_001035232 |
| h$Na_V$1.3 | RLMTQDYWEN | RVLCGEWIET | VATFKGWMDI | ITTSAGWDGL | NP_008853 |
| h$Na_V$1.4 | RLMTQDYWEN | RILCGEWIET | VATFKGWMDI | ITTSAGWDGL | NP_000325 |
| h$Na_V$1.5 | RLMTQCFWER | RILCGEWIET | VATFKGWMDI | ITTSAGWDGL | NP_932173 |
| h$Na_V$1.6 | RLMTQDFWEN | RVLCGEWIET | VATFKGWMDI | ITTSAGWDGL | NP_055006 |
| h$Na_V$1.8 | RLMTQDSWER | RILCGEWIEN | VATFKGWMDI | ITTSAGWDGL | NP_006505 |
| h$Na_V$1.9 | RLMTQDSWEK | RILCGEWIEN | VATFKGWMDI | ITTSAGWDSL | NP_054858 |

**Supplemental Table 2:** ST-2262 $IC_{50}$ against $Na_V1.1 – 1.8$, determined by manual patch clamp electrophysiology

| Isoform | $IC_{50}$ (μM) | 95% CI | # of data points |
|---|---|---|---|
| $Na_V1.1$ | >100 | | 6 |
| $Na_V1.2$ | >100 | | 5 |
| $Na_V1.3$ | 65.3 | 62.7–68.1 | 7 |
| $Na_V1.4$ | 80.7 | 71.1–93.3 | 35 |
| $Na_V1.5$ | >100 | | 18 |
| $Na_V1.6$ | 17.9 | 14.8–22.1 | 30 |
| $Na_V1.7$ | 0.072 | 0.064–0.082 | 70 |
| $Na_V1.8$ | >100 | | 3 |

Note: 100 μM was the highest concentration tested

**Supplemental Table 3:** ST-2262 $IC_{50}$ against $Na_V1.1$ – 1.7, conducted by an independent lab using the PatchXpress automated electrophysiology platform

| Isoform | $IC_{50}$ (µM) | 95% CI | # of data points |
|---|---|---|---|
| $Na_V1.1$ | >100 | | 9 |
| $Na_V1.2$ | >100 | | 17 |
| $Na_V1.3$ | >100 | | 11 |
| $Na_V1.4$ | >100 | | 20 |
| $Na_V1.5$ | >100 | | 17 |
| $Na_V1.6$ | 25.8 | 21.7–30.8 | 18 |
| $Na_V1.7$ | 0.057 | 0.051–0.064 | 24 |

Note: 100 µM was the highest concentration tested

**Supplemental Table 4:** ST-2262 $IC_{50}$ against $Na_V1.7$ with different protocols determined by manual patch clamp electrophysiology

| Protocol | $IC_{50}$ (μM) | 95% CI | # of data points |
|---:|:---:|:---:|:---:|
| Resting State* | 0.123 | 0.104–0.145 | 17 |
| State-dependent | 0.087 | 0.056–0.120 | 6 |
| Use-dependent | 0.112 | 0.015–0.357 | 5 |

*Resting state recordings were only included if the ST-2262 lot was also tested on state- and use-dependent protocols.

**Supplemental Table 5:** ST-2262 $IC_{50}$ against $Na_V1.7$ from multiple species using the PatchXpress automated electrophysiology platform

| $Na_V1.7$ Species | $IC_{50}$ (µM) | 95% CI | # of Data points |
|---:|:---:|:---:|:---:|
| Cynomolgus Macaque | 0.101 | 0.073–0.140 | 10 |
| Mouse | 3.78 | 3.23–4.43 | 18 |
| Rat | 4.95 | 4.17–5.87 | 13 |

**Supplemental Table 6:** ST-2262 $IC_{50}$ against mutant and WT mouse and human $Na_V1.7$ channels determined by manual patch clamp electrophysiology

| Isoform | $IC_{50}$ (µM) | 95% CI | # of data points |
|---|---|---|---|
| m$Na_V1.7$ WT | 2.57 | 2.30–2.87 | 9 |
| m$Na_V1.7$ M1407T/D1408I | 0.130 | 0.055–0.307† | 4 |
| h$Na_V1.7$ WT | 0.039 | 0.032–0.047 | 5 |
| h$Na_V1.7$ D1690N | >100 | | 7 |

†Asymptotic 95% CI calculated

**Supplemental Table 7:** ST-2262 Pharmacokinetics– IV administration in cynomolgus monkey[a]

| | |
|---|---|
| **$t_{1/2}$ (h)** | 3.50 ± 0.013 |
| **$Vd_{ss}$ (L/kg)** | 0.27 ± 0.021 |
| **Cl (ml/min/kg)** | 1.17 ± 0.037 |

[a]0.5 mg/kg ST-2262 administered. Values are the mean ± SD (n=3)

### Characterization data for guanidinium toxin analogues

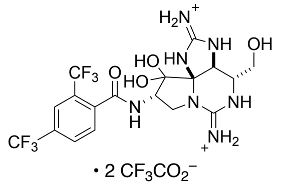

**ST-282.** $^{1}H$ NMR ($D_2O$, 400 MHz) $\delta$ 8.08 (s, 3H), 4.99 (dd, 1H, $J$ = 9.2, 9.2, Hz) 4.84 (s, 1H), 4.24 (dd, 1H, $J$ = 9.6, 9.6 Hz), 3.74–3.65 (m, 3H), 3.38 (dd, 1H, $J$ = 9.2, 9.2 Hz) ppm. LRMS (ES+, m/z) calcd for $(M+H)^+$ $C_{18}H_{20}F_6N_7O_4^+$: 512.4; found: 512.2.

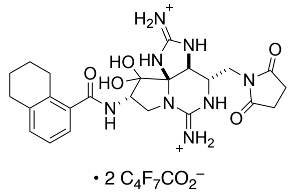

**ST-2262.** $^{1}H$ NMR ($D_2O$, 400 MHz) $\delta$ 7.42–7.30 (m, 3H), 5.07–5.02 (m, 1H), 4.98 (s, 1H), 4.34 (t, $J$ = 9.8 Hz, 1H), 4.11–4.02 (m, 1H), 3.96–3.91 (m, 1H), 3.67 (dd, $J$ = 14.0, 3.4 Hz, 1H), 3.57 (t, $J$ = 9.6 Hz, 1H), 2.94 (m, 8H), 1.93–1.88 (m, 4H) ppm; $^{13}C$ NMR ($D_2O$, 100 MHz) $\delta$ 181.0, 174.4, 157.4, 155.3, 138.8, 135.0, 134.5, 131.5, 125.5, 124.1, 97.7, 82.1, 58.5, 51.8, 51.0, 46.1, 38.8, 29.1, 28.0, 26.4, 22.4, 22.2 ppm. LRMS (ES+, m/z) calcd for $(M+H)^+$ $C_{24}H_{31}N_8O_5^+$: 511.6; found: 511.3.

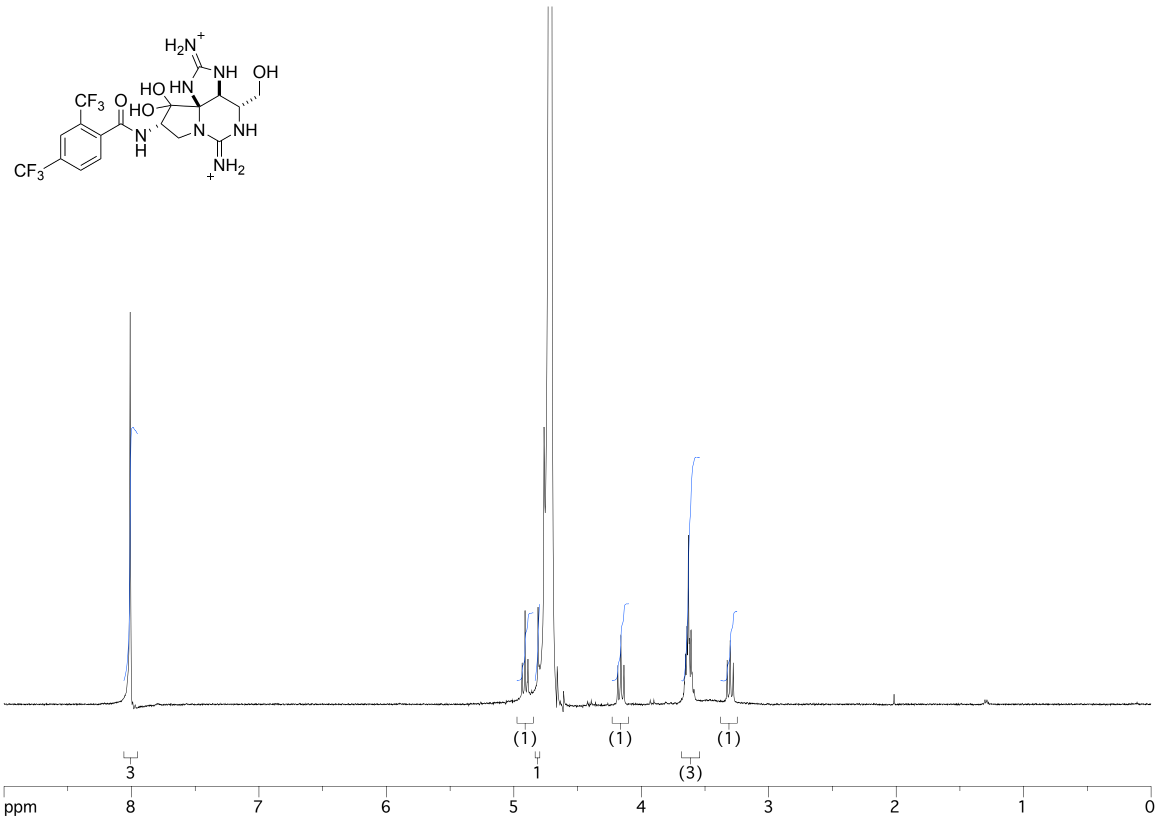

$H_2N^+$
NH
OH
HN
HO
HO
$CF_3$
O
N
H
N
NH
$NH_2$
$CF_3$
(1)
(1)
(1)
3
1
(3)
ppm
8
7
6
5
4
3
2
1
0


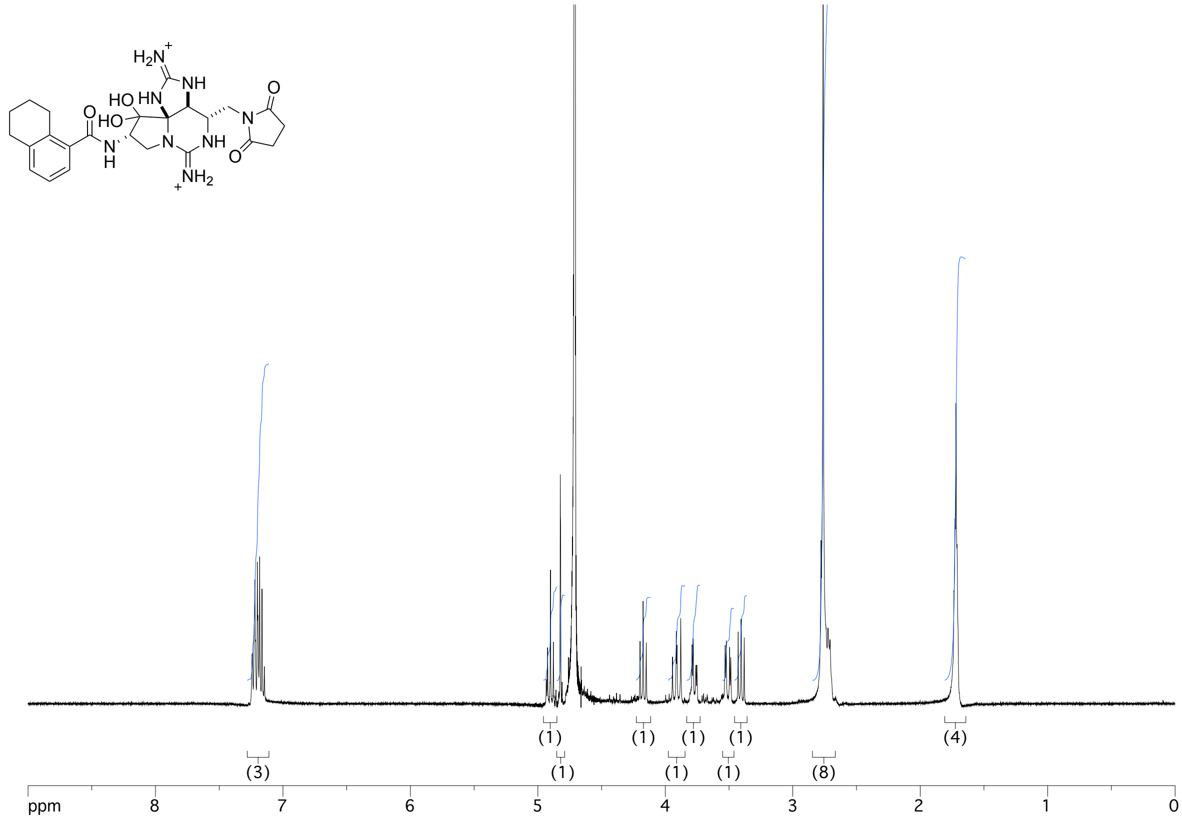

$H_2N^+$
NH
HO
HN
O
HO
N
H
N
NH
O
N
O
$NH_2$
(1)
(1)
(1)
(1)
(4)
(3)
(1)
(1)
(1)
(8)
ppm
8
7
6
5
4
3
2
1
0
